# Fishing Cat Cell Biobanking for Conservation

**DOI:** 10.1101/2022.07.01.498384

**Authors:** Woranop Sukparangsi, Ampika Thongphakdee, Santhita Karoon, Nattakorn Suban Na Ayuttaya, Intira Hengkhunthod, Rachapon Parkongkeaw, Rungnapa Bootsri, Wiewaree Sikaeo

**Affiliations:** Department of Biology, Faculty of Science, Burapha University, Chon Buri, Thailand; Wildlife Reproductive Innovation Center, Research Department, Bureau of Conservation and Research, Zoological Park Organization under the Royal Patronage of H.M. the King, Bangkok, Thailand

**Keywords:** cryopreservation, biobank, fishing cat, wild felid, fibroblast, reprogramming

## Abstract

Establishment of biobank to keep wildlife cells secure long-term conservation. Fishing cat (*Prionailurus viverrinus*) is one of Vulnerable wild felids, currently under threaten by wetland destruction and other human activities. Here we aimed to generate cell biobanking of fishing cats by deriving various sources of primary cells from the living and postmortem animals and enhancing their expandable potency by virus-free cellular reprogramming. We show that cells can be propagated from several tissues harvested from both living and dead fishing cats with different derivation efficiency. Testes from the postmortem animals contain several tissues that can be derived primary cells as well as putative alkaline phosphatase positive and SOX2 positive adult spermatogonial stem cells. Primary cells from ear pinna and abdomen sources can only be obtained from the living fishing cats. These primary cells exhibited sign of cell senescence after a few sub-cultures, limited its usability for downstream applications. This obstacle can be overcome by reprogramming via either nucleofection or liposome-based DNA/RNA delivery. The putative iPSC colonies as well as expandable induced cells from episomal-based reprogramming appeared to be a suitable choice for expansion of cells for cryopreservation. Thus, here we provide current conservation plan using cell technology for fishing cats and also recommendation of tissue collection and culture procedures for zoo researches to facilitate the preservation of cells from postmortem animals and living animals.

**Highlight:** Biobanking of viable cells is essential to provide long-term security of wildlife existence. Current cell technology enables us to cultivate primary cells and adult germ cells from tissues of living and postmortem fishing cats for cryopreservation. The primary cells exhibited limited proliferation and cell senescence, which can be overcome by reprogramming the somatic cells toward pluripotent state. Here we explored the challenge of tissue collection from fishing cat and several virus-free approaches to induce cellular reprogramming in the fishing cat cells and provided insight into the techniques and conditions to enhance cell expansion, which support the success of generation of fishing cat cell biobank.

## Introduction

Distribution records of wild felid in Thailand via camera-trap surveys, radio-telemetry studies, direct sightings, interviews as well as searches for signs from several efforts show the presence of seven small to medium cat species including Asiatic golden cat *Catopuma temminckii*, jungle cat *Felis chaus*, mainland clouded leopard *Neofelis nebulosa*, marbled cat *Pardofelis marmorata*, leopard cat *Prionailurus bengalensis*, flat-headed cat *Prionailurus planiceps*, and fishing cat *Prionailurus viverrinus* [1-4], and two large cat species including Indochinese tiger *Panthera tigris corbetti* and leopard or panther *Panthera pardus* [2,5-6].

Fishing cat plays important roles in wetland ecosystem; however, in recent years, their wetland habitats are destroyed by human threats such as prawn farming industry, illegally established aquaculture ponds, human settlement, excessive hunting, deforestation, depletion of fish stocks from over-fishing, and incidental poisoning [7-9]. The fishing cat is now included on “Vulnerable” status (IUCN Red List of Threatened Species) [10] and CITES Appendix II and protected by national legislation. Fishing cats in captivity at Zoological Park Organization of Thailand are currently under conservation programs aimed to improve its breeding and produce more offspring, which can be eventually re-introduced back to natural habitats. Rescuing of endangered or vulnerable wildlife animals by breeding program at the zoo involves assisted reproductive technology (ART), which become urgent trends to secure good genetics for long-term conservation. The storage of valuable genetics of wild animals can be performed in several ways including cryopreservation of gametes (spermatozoa and oocyte) and embryos for future *in vitro* fertilization (IVF), artificial insemination (AI) and embryo transfer [11-13]. To date, Santymire et al., 2011 and Thongphakdee et al., 2018 [14-15] provided better understanding of reproductive biology in fishing cat and demonstrated a potential application in its captive breeding. However, there are some restrictions for developing ART in fishing cat including restricted number of fertile fishing cats, high variability of ovarian response to estrus and ovulation induction for AI [15] and low fertilization success [16].

Another approach to secure good genetics of fishing cats as well as other wild animals is to establish primary cells, in particular fibroblasts with some restricted degree of propagation and stem cell lines such as embryonic stem cells (ESCs), mesenchymal stem cells (MSCs) and induced pluripotent stem cells (iPSCs) with more capacity to self-renewal and differentiation. The preservation of those cells in liquid nitrogen-containing cryotank generates invaluable wildlife biobank, or “Frozen Zoo” for future conservation approaches. *In vitro* culture of various cell types also provides great benefit of reducing and replacement of using animals for experimentation in 3Rs (Reduction, Refinement and Replacement) model [17].

Tissues in primary culture can be harvested from either live animals or freshly dead/post-mortem animals [18-20]. Fibroblast culture has already been achieved among several mammalian species, mostly in mouse (*Mus musculus*) and human and in other mammals such as horse, dog, drill, rhinoceros, and elephant [20-23]. In felid species, fibroblasts were derived in domestic cat for iPSC generation and intraspecific feeder cells to support induction of iPSC and cat ESC derivation and in wild cats including Bengal tiger, jaguar, serval and snow leopard for iPSC generation [24-27]. Hence, in this study we aimed to establish cell biobanking for fishing cats to preserve its genetics for long-term conservation. During a decade of fishing cat projects in our zoo organization, here we reported the successful attempts to derive fibroblasts from different sources including living and dead fishing cats in captivity and from natural resources. Here we also examined the usability of our fibroblasts for cellular reprogramming and provided the insight into current challenges to these approaches.

## Methods

### Animal Ethics

This project was permitted to perform under project “Development of fundamental science and innovation for sustainable fishing cat conservation” with sub-project “Conservation of fishing cat (*Prionailurus viverrinus*) by using innovation of stem cell technology” by wild animal ethics committee of Zoological Park Organization of Thailand under the Royal Patronage of H.M. the King (Protocol number: 630960000030 granted to PI of the project-A.T.). Permission to work with wild fishing cats in the Natural Parks of Thailand was granted by Department of National Park Wildlife and Plant Conservation-Thailand (Permission number: Tor Sor (in Thai) 0907.4/17939 (Issued date: 22/09/2021). Tissue collection from postmortem fishing cats and during artificial insemination were performed by veterinarians at Animal Hospital Unit, Khao Kheow Open Zoo (KKOZ), Zoological Park Organization, Chon Buri, Thailand. Cell culture was performed at Tissue Culture Facility at Wildlife Reproductive Innovation Center (WRIC), Research Department, Bureau of Conservation and Research, KKOZ and genetic materials and chemicals in this study were used under Certificate of Biosafety (22/2559) approved by Biosafety Committee, Burapha University.

### Primary culture and cryopreservation

Collected tissues (skin and testis) were washed with Dulbecco’s Phosphate-Buffered Saline (DPBS, ThermoFisher) containing Penicillin (10000 Unit/mL)-Streptomycin (10 mg/mL) (PS) solution (Sartorius) and 500 µL 10X Amphotericin B (AmB) Solution (Sartorius) for 3 times. Removal of subcutaneous adipose tissues and intact hairs (for skin tissues) was done to avoid bacterial and fungal contamination and presence of lipid droplets. To activate the outgrowth of fibroblasts from the tissues using wound healing process, several small cuts were made using scalpel and the tissues were cut into approximately 2×2 mm. The excised tissues were then placed (3-4 pieces) on gelatin (Attachment Factor, ThermoFisher)-coated dish (35 mm) or 12 well plates already for 10 min for better explant attachment. Then warm complete fibroblast medium was added to the dish/plate as lowest volume as possible to avoid tissue floating and ensure the outgrowth. The primary culture was placed in the CO2 incubator (5% CO2 in humidified 95% air at 37 °C). To avoid contamination, the cultures were monitored every day and medium changes should be done every day without disturbing the explant attachment. The complete fibroblast medium (50 mL) was composed of Dulbecco’s Modified Eagle Medium (DMEM) high-glucose medium (ThermoFisher and Sartorius), MEM Non-essential Amino Acid Solution (NEAA, Sigma, Merck), Glutamax (ThermoFisher), Sodium Pyruvate (Sigma, Merck) and antibiotics PS-antimycotics AmB (Sartorius), unless stated otherwise. For cryopreservation, the cells were resuspended in freezing medium containing complete fibroblast medium with 10% dimethyl sulfoxide (DMSO) or Recovery™ Cell Culture Freezing Medium (ThermoFisher) and placed in - 80 °C overnight using Mr.Frosty™ Freezing Container and next day transferred to liquid nitrogen for long-term storage. All cells in this study are currently stored in Biobank Liquid Nitrogen Tank Facility under Zoological Park Organization (ZPO) of Thailand. Further detail of cell storage and material transfer is available upon request to correspondence (A.T. or ZPO Research Bureau).

### MTS assay

MTS colorimetric assay kit (Abcam) was used to quantify cell viability of transfected cells. For nucleofection, the transfected cells were seeded into 96-well plate (10,000 cells/well) in the presence or absence of RevitaCell™ supplement (Gibco). RevitaCell™ supplement was added at final concentration at 0.5X. At day 4 post nucleofection, fresh fibroblast medium was changed prior performing MTS assay (200 uL/well). Then 20 uL MTS solution was added and incubate for 4 hours at 37 °C. Absorbance at 490 nm was measured by microplate reader (Thermo Scientific™ Multiskan™ GO Microplate Spectrophotometer) with Skanlt™ Software. Four independent experiments with three technical replicates each were performed for measuring cell viability.

### Alkaline phosphatase live staining

To identify pluripotent stem cells before cryopreservation for future uses, we used Fluorescein-based Alkaline Phosphatase (AP) Live Stain (ThermoFisher) to detect activity of AP in cell extracts from seminiferous tubules and epididymis. Briefly, we prepared diluted AP stain (1:500) and Hoechst33324 mixed with medium and stained the cells for 20 minutes before imaging with fluorescent microscope.

### Immunofluorescence

To observe the presence of Spermatogonial Stem Cells (SSC), extracts from seminiferous tubules from the fishing cat were fixed with 4% paraformaldehyde (PFA) for 15 mins at room temperature and washed with DPBS for 3 times. In order to permeabilized the plasma membrane, the fixed cells were treated with 0.1% Triton X-100 in PBS for 15 mins at room temperature and washed with DPBS for 3 times with each time incubated for 5 mins. Then the cells were treated with 1% BSA in DPBS for 1 hour at room temperature. The cells were treated with primary antibodies SOX2 (AB5603, Merck), at dilution 1:300, for over-night at 4 °C and washed with DPBS 3 times next day before secondary antibody staining. On the next day, the cells were stained with Alexa Fluor 647 (ThermoFisher, 1:800) and Hoechst33342 (ThermoFisher, 1:200) in dark for 1 hr at room temperature and washed with DPBS 3 times. Fluorescent micrographs were taken by inverted microscope Eclipse Ti-S Inverted Research Microscope (Nikon) and digital camera.

### Nucleofection

Nucleofection™ programs in 4D Nucleofector™ system included CA-137 (specific for human iPSCs and mammalian fibroblast with Primary Cell 2 (P2) and 3 (P3) solution kit), DS-150, EH-100, EN-150, EO-114 (specific for mammalian fibroblast recommended by Lonza and compatible with P2 solution kit), FF-135 (specific for human fibroblast and dental palp cells [28-30]) and DT-130 (specific for neonatal Normal Human Dermal Fibroblast, NHDF-neo cell lines compatible with P2 solution and human fibroblast [31]). Small scale of 4D-NucleoFector™ kits (P2 Primary Cell 4D-Nucleofector® X Kit S 32 RCT, V4XP-2032), composed of 16-well Nucleocuvette™ Strip and Primary Cell 2 (P2) solution, were used for nucleofection. To prepare fibroblast cell transfection with small scale, the cells at 70% confluency were subcultured, counted by hemocytometer, transferred 10^5^ cells to new microcentrifuge tube, and then centrifuged at 90g for 5 minutes, supernatant was aspirated supernatant and cell pellet were gently resuspended in 20 µL of P2 solution, 400 ng of pmaxGFP™ vector was added and gently mixed, transferred cell suspension into Nucleocuvette™ Strip, nucleofected with programs of interest, kept the transfected cell suspension at room temperature for 10 minutes, 80 µL of complete fibroblast medium (CF) added to Nucleocuvette™, transferred transfected cells into non-coated 4-well plates containing 420 µL complete fibroblast medium per well, and incubating cells in a CO2 incubator. GFP expression was observed under fluorescence microscope.

### Senescence test

Senescence β-Galactosidase Staining (Abcam) was used to detect sign of cell senescence according to manufacturer’s instruction. Briefly, the fibroblasts were washed once with DPBS, fixed with fixative solution for 10 minutes and then washed with DPBS twice. The cells were then stained with staining solution mix containing Solution A, Solution B and X-gal for overnight at 37 °C. On the next day, the stained cells were stored in 70% Glycerol before imaging with an inverted microscope.

### Flow cytometric analysis

To monitor transfection efficiency, GFP expression was detected by flow cytometry (FlowSight® Imaging Flow Cytometer, Luminex). At day 4 post nucleofection, transfected cells were washed once with DPBS, dissociated with 0.25% Trypsin-EDTA for 3 minutes, neutralizing with fibroblast medium, spinning down cell pellet and resuspending in DPBS. Hoechst33342 (1 µg/mL) were then added to cell suspension to stain all cells. Flow cytometric data were analyzed using FCS Express 7 software. GFP/Hoechst33342 cell populations were gated and transfection efficiency was calculated from percentage of double positive cell populations with GFP and Hoechst33342 against total Hoechst33342 positive cells. Four independent experiments were performed for measuring transfection efficiency.

### Reprogramming assay

PiggyBAC transposon (PB) vector MKOS-mOrange (Gifted by Dr.Keisuke Kaji) contains 4 mouse reprogramming factors: C-Myc, Klf4, Oct4 and Sox2 under CAG-promoter and mOrange reporter. The DNA vector was transfected by nucleofection. The transfected cells were cultured with iPSC induction medium containing Advanced DMEM (ThermoFisher), 10% FBS, MEM Non-essential Amino Acid Solution (NEAA, Sigma, Merck), Glutamax (ThermoFisher), Sodium Pyruvate (Sigma, Merck) and antibiotics PS-antimycotics AmB (Sartorius) with 10 ng/mL human Leukemia Inhibitory Factor (hLIF, Peprotech), unless stated otherwise. The cells were re-seeded onto irradiated MEFs or other coating matrixes including Attachment Factor/gelatin (ThermoFisher), Geltrex (ThermoFisher) and vitronectin (ThermoFisher). PB vector pC6F (addgene: 140826 [32]) contains Tet-on system regulating the expression of polycistronic cassettes of 6 reprogramming factors: human OCT4, SOX2, KLF4, CMYC, KLF2 and NANOG and TdTomato reporter [32]. We transfected pC6F alongside with transposase vector (pCy43, Sanger Institute) and PB rtTA vector (addgene 126034) using Lipofectamine 3000 (ThermoFisher) for overnight. The transfection was performed for one time and on day 3, we selected cells carrying pC6F construct with puromycin (2 ug/mL) and induced the expression of reprogramming factors with Doxycycline in the iPSC induction medium. Episomal vector system is composed of pCXLE-hSK (vector with SOX2 and KLF4; addgene: 27078), pCXLE-hUL (vector with L-MYC and LIN28; addgene: 27080), pCXLE-hOCT3/4 (vector with OCT3/4; addgene: 27076), pCXWB-EBNA1 (vector with EBNA1; addgene: 37624). These vectors were nucleofected into the cells. The transfected cells were cultured in the iPSC medium, unless state otherwised. Commercial media were also tested for reprogramming, including NutriStem hPSC XF Medium (Sartorius), medium containing Knockout Serum Replacement (KOSR, ThermoFisher) and Essential 8 medium (ThermoFisher). For self-replicating RNA (srRNA) system, we generated RNA from T7-VEE-OKSiM, T7-VEE-OKSiG, T7-VEE-GFP plasmids using *in vitro* transcription technique with HiScribe™ T7 quick high yield RNA synthesis kit, according to manufacturer’s protocol. Synthesized RNA was modified by Vaccinia Capping System (NEB), mRNA cap 2’-O-methyltransferase (NEB) and *E. coli* Poly(A) Polymerase (NEB). The fibroblasts were transfected with srRNA using Lipofectamine™ MessengerMax™ (ThermoFisher) according to adjusted Yoshioka & Dowdy method [33]. The transfected cells were cultured in iPSC medium with 200 ng/mL of recombinant viral B18R protein (R&D Systems). B18R was removal once the transfected cells were ready to re-seed onto irradiated MEFs.

### Statistical analysis

Data from three to four independent experiments (with at least three technical replicates) are presented as the mean±SD (Standard Deviation). Statistical analyses were performed using Student’s t test to determine statistical significance between the groups.

## Results

### Adult dermal fibroblast culture from abdominal dermis of living fishing cats

Skin biopsy to collect dermis for producing fibroblast culture, an available source for cellular reprogramming, from living fishing cats have been limited due to invasive procedures. Within assisted reproductive technology (ART) program of the zoo, direct harvest of the dermis in the abdominal skin of fishing cat during artificial insemination (AI) operation were conducted (Figure 1Ai-iii), enabling us to derive dermal fibroblasts from a female fishing cat. As shown in Figure 1B, outgrowth of the fibroblasts emerged around the edge of the tissue explants within five days. Cell with epithelial character was not detected, indicating the absence of keratinocytes (Figure 1B-E). The dermal fibroblasts in the primary culture expanded to reach its 90% confluency at day 16 with homogenous fibroblastic morphology (Figure 1D). However, as the collection of tissue from abdominal dermis was closed to subcutaneous layer containing rich adipose tissue, lipid droplets were found during culture (Figure 1E). Secondary culture was done by removing explant tissues and sub-cultured the fibroblasts into appropriate culture vessels (split ratio of 1:2; Cell density at 2.0 × 10^4^ cells/cm^2^). Fibroblasts expanded to reach its 90% confluency within 4 days after the first passaging (Figure 1F). It is noteworthy that seeding the fibroblasts of fishing cat at too low cell density generally led to less cell expansion. In early passages, the fibroblasts exhibited spindle shape (Figure 1F) while at later time the cell area expanded in less nucleus:cytoplasm ratio manner (Figure 1G). At the late passages, the fibroblast culture was deficient due to longer trypsinization time needed for subculture and less cell expansion. By removing supplements and measuring cell division rate with MTS assay, we also show the essential components of the fibroblast medium to support cell growth and division. As shown in Figure 1H, removing either sodium pyruvate or non-essential amino acids or both did not reduce cell proliferation while removing Glutamax caused a significant reduction of cell proliferation. The effect became prominent from the complete removal of all supplements. Thus, additional supplements did help to improve fibroblast cell expansion. In addition, increasing percent of fetal bovine serum (FBS) from 10% to 20% did not significantly enhance cell proliferation (Figure 1H). To improve the fibroblast culture, we next tested addition of basic Fibroblast Growth Factor (bFGF or FGF2) and found that bFGF enhanced proliferation of the fibroblasts (Figure 1I-K).

**Figure 1.**
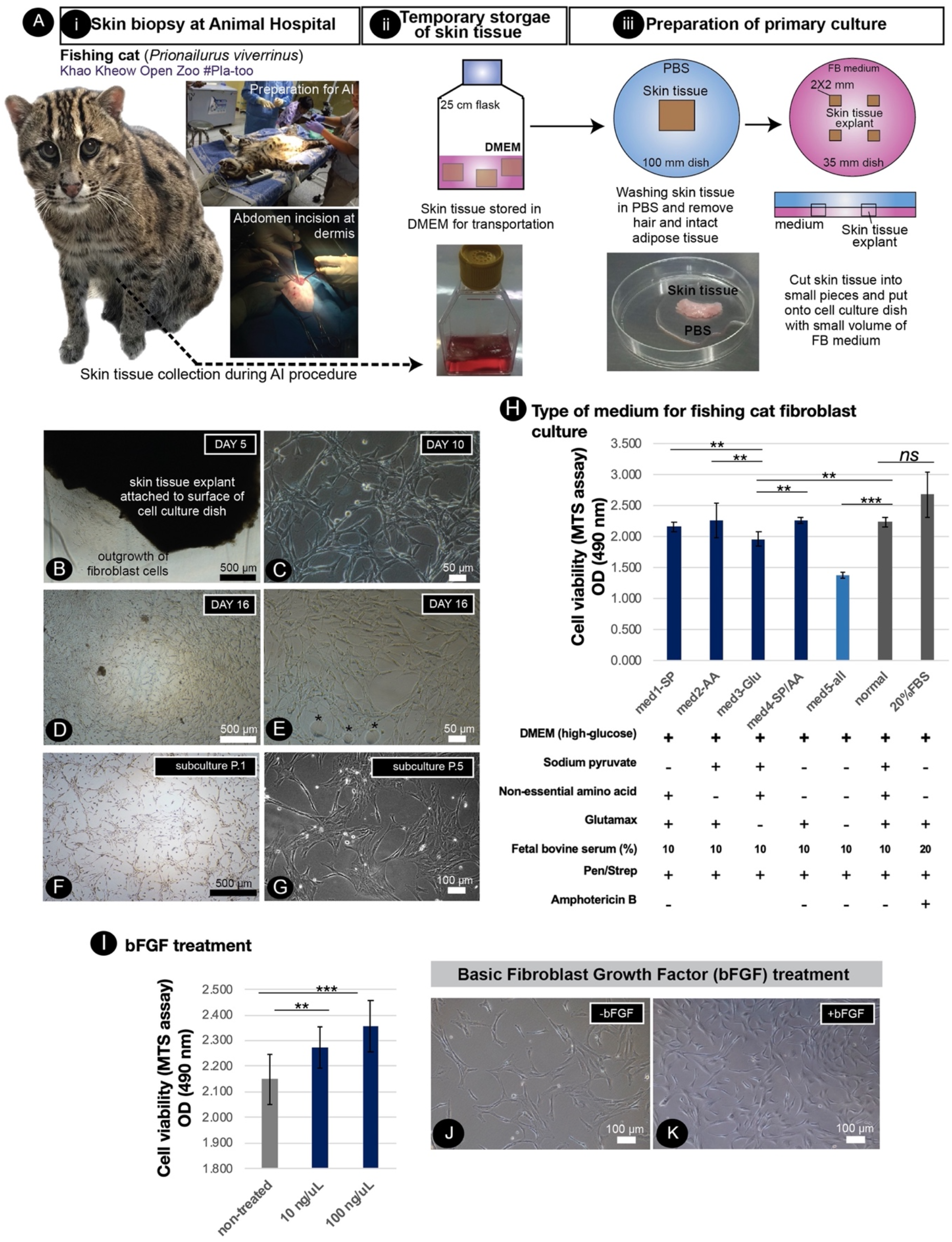
Primary culture of fishing cat cells and conditions supporting fishing cat cell expansion. A) Tissue collection and preparation for primary culture i) Skin biopsy was performed by veterinarians during artificial insemination of fishing cat. ii) The skin tissues were then transported to Tissue Culture Facility in DMEM or complete fibroblast (FB) medium with antibiotics. iii) The skin tissues were washed with PBS and dissected into small pieces and the tissue explants were cultured in FB medium. B) The adult fibroblasts were expanded from skin tissue explant as early as day 5. C) Primary culture shows morphology of fishing cat fibroblast at day 10 D) Fibroblasts expanded and reached 80% confluency within 16 days. E) Primary culture contained some oil droplets (*) due to high subcutaneous fat connected to the dermis layer. F)-G) Culture of the adult dermal fibroblasts after subculture from the primary culture. H) Adult dermal fibroblasts from fishing cat at passage 2-4 were cultured in different medium conditions. Bar graph shows cell viability of fibroblasts under different medium conditions at day 7 post treatment. The cell viability was measured by MTS assay using spectrophotometer (at 490 nm). I) The fibroblasts were treated with basic fibroblast growth factor (bFGF) at final concentration 10 ng/µL and 100 ng/µL. Bar graph shows MTS assay-based cell viability of fibroblasts at day 4 post bFGF treatment. J) Brightfield photograph shows morphology of fibroblasts without bFGF. K) Brightfield photograph shows morphology of fibroblasts with bFGF at day 2 post treatment. Asterisks (** and ***) indicate significant difference (p<0.05 and p<0.01 respectively, Student’s t test) and “ns” indicates not significant (p<0.05).

### Adult dermal fibroblast culture from ear pinna of living fishing cats

To preserve more genetic variation of fishing cats, cryopreservation of cell samples collected from natural resources is required. Under permission of National Park of Thailand (in Animal Ethics section), two fishing cats from nature were caught to collect samples including skin biopsy. Due to nature of fishing cats in prey capture by swimming, we collected small skin tissues from the ear pinna (Figure 2A,F), not other parts of abdominal and leg area, of anesthetized fishing cats by veterinarian team. The skin tissue (Figure 2A, F) was kept cold (4°C) in complete fibroblast medium during transportation to Tissue Culture Facility. The tissue explants (Figure 2B, G) were cultured within 4-5 hours after the skin biopsy. Primary culture of freshly collected ear pinna tissue showed that the fibroblast outgrowth can be observed within three days from most of explant tissues (Figure 2C, D, H, I). However, epithelial-like keratinocytes were also expanded in advanced DMEM (AD) medium (Figure 2D right, I right), but not in the fibroblast (FB) medium (Figure 2C, H), which the proportion of keratinocytes/fibroblasts diminished after passaging in DMEM-based medium (Figure 2E, J). The tissue explants from living fishing cat can be re-explanted for at least three times and producing fibroblast culture sufficient to freeze for 30 cryovials (approximately 100,000 cells/vial), Table 1. It is noteworthy that cell cultures of tissues collected from natural resources caused regular fungal contamination, even the presence of Amphotericin B. We also compared duration of medium change and found that primary culture with medium change everyday produced more fibroblasts while medium change every 2/3-day period caused 100% contamination, shown as overall result in Figure 2.

**Figure 2.**
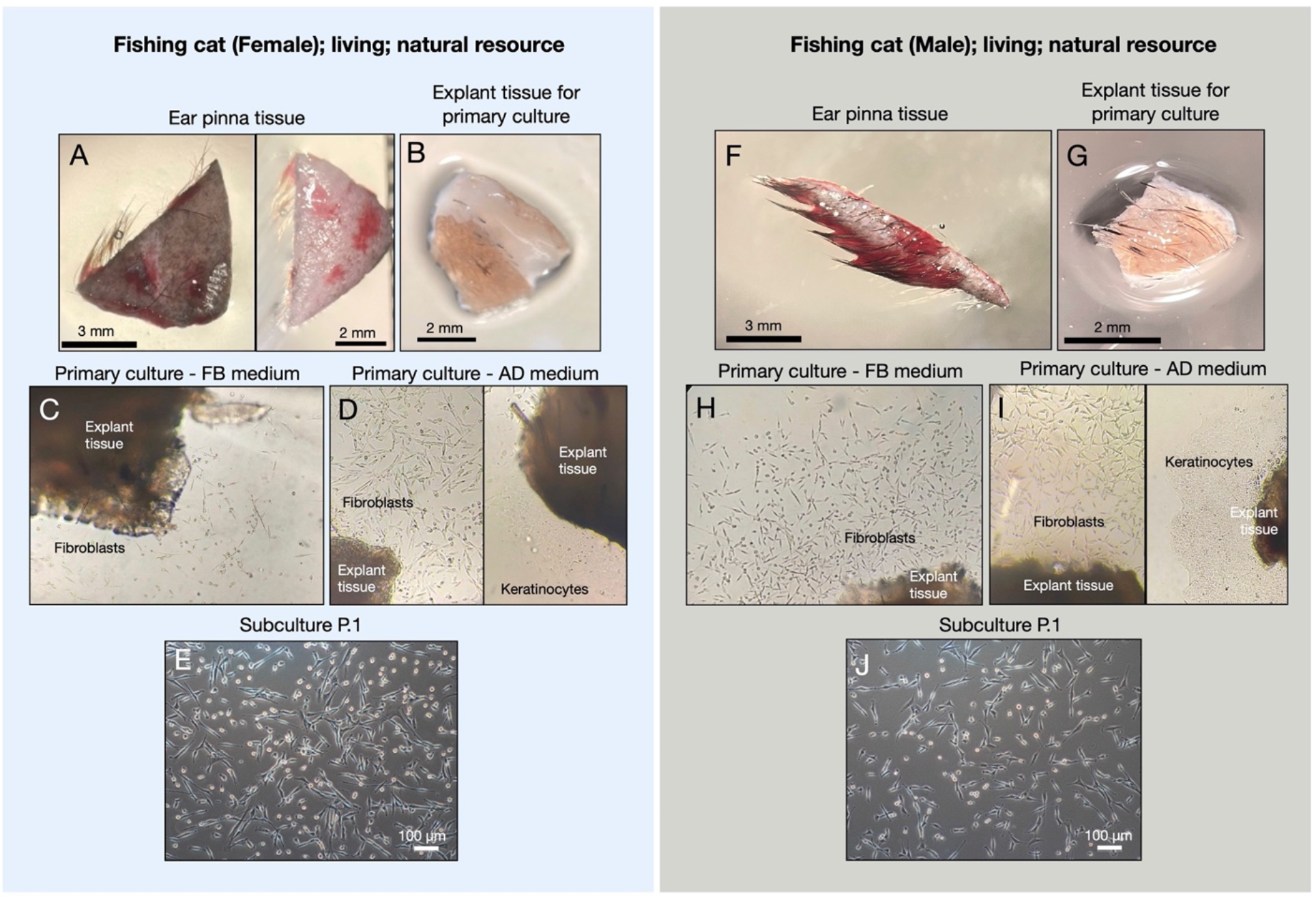
Primary culture of fishing cat tissues collected from natural resource. Ear pinna tissues from female fishing cat (A) and male fishing cat (F) were collected and prepared for primary culture. The collected tissues were washed and dissected into small pieces (explant) before attaching on gelatin-coated culture dishes (B, G). Th explants were cultured in complete fibroblast (FB) medium and Advanced DMEM (AD) medium (C-D, H-I). Fibroblast outgrowth was found in both FB and AD medium (C-D, H-I) while Keratinocyte outgrowth was found only in AD medium (D, I). E, J) Subculture at passage 1 (P.1) after primary culture reaching 90% confluency.

**Table 1.** Primary culture of fishing cat tissues from different sources

| Sources for tissue collection | Sex | Estimated age | Live or Postmortem animals | Days of the first appearance of fibroblast outgrowth | Number of re-explant | Number of vials for cryopreservation |
| --- | --- | --- | --- | --- | --- | --- |
| *Natural resources* |  |  |  |  |  |  |
| Ear pinna | Male | Young adult | Live | 2-3 days | 3-4 times | 19 |
| Ear pinna | Female | Young adult | Live | 2-3 days | 3-4 times | 11 |
| *Captivity* |  |  |  |  |  |  |
| Abdominal tissues (AI) | Female | Adult | Live | 15 days | 0 time | 7 |
| Abdominal tissues (AI) | Male | Adult | Live | 13 days | 0 time | 3 |
| Ear pinna | Female | Adult | Postmortem | 13 days | 0 time | 0 |
| Ear pinna | Female | Baby | Postmortem | 5 days | 0 time | 2 |
| Testis | Male | Adult | Postmortem | 3 days | 2 times | 5 |

### Primary culture of tissues from postmortem fishing cats

From a postmortem fishing cats at 3 months old (3FC) and 11 years old (11FC), we can retrieve two parts of the body including ear pinna (both 3FC and 11FC) and testes (11FC). Explants from the ear pinna were seeded after one day postmortem. We can derive fibroblasts from ear pinna only from 3FC (Figure 3A-C) but cannot obtain any primary cells after prolonged culture for a month from 11FC (Figure 3D). For tissue collection from testes, the fibroblasts can be obtained from testicular wall and epididymis (Figure 3E, F). Comparing the testicular fibroblasts between P.1 and P.4, we observed stronger signal of senescence in P.4 (Figure 3G, H). This indicates that the usability of the adult fibroblasts from testis is limited to only early passages. We also examined the extraction of cells from seminiferous tubules and epididymis to observe the presence of spermatogonial stem cells (SSCs). Using immunofluorescence with an antibody to detect SOX2 protein, we found SOX2-positive cells in the culture from seminiferous tubules, indicating the plausible presence of SSCs (Figure 3I). We also found alkaline phosphatase positive cells within the seminiferous tubule but not in the epididymis (Figure 3J). Thus, from postmortem fishing cat, we can cryopreserve cells from various sources including testicular wall, epididymis and putative SSCs from seminiferous tubules.

**Figure 3.**
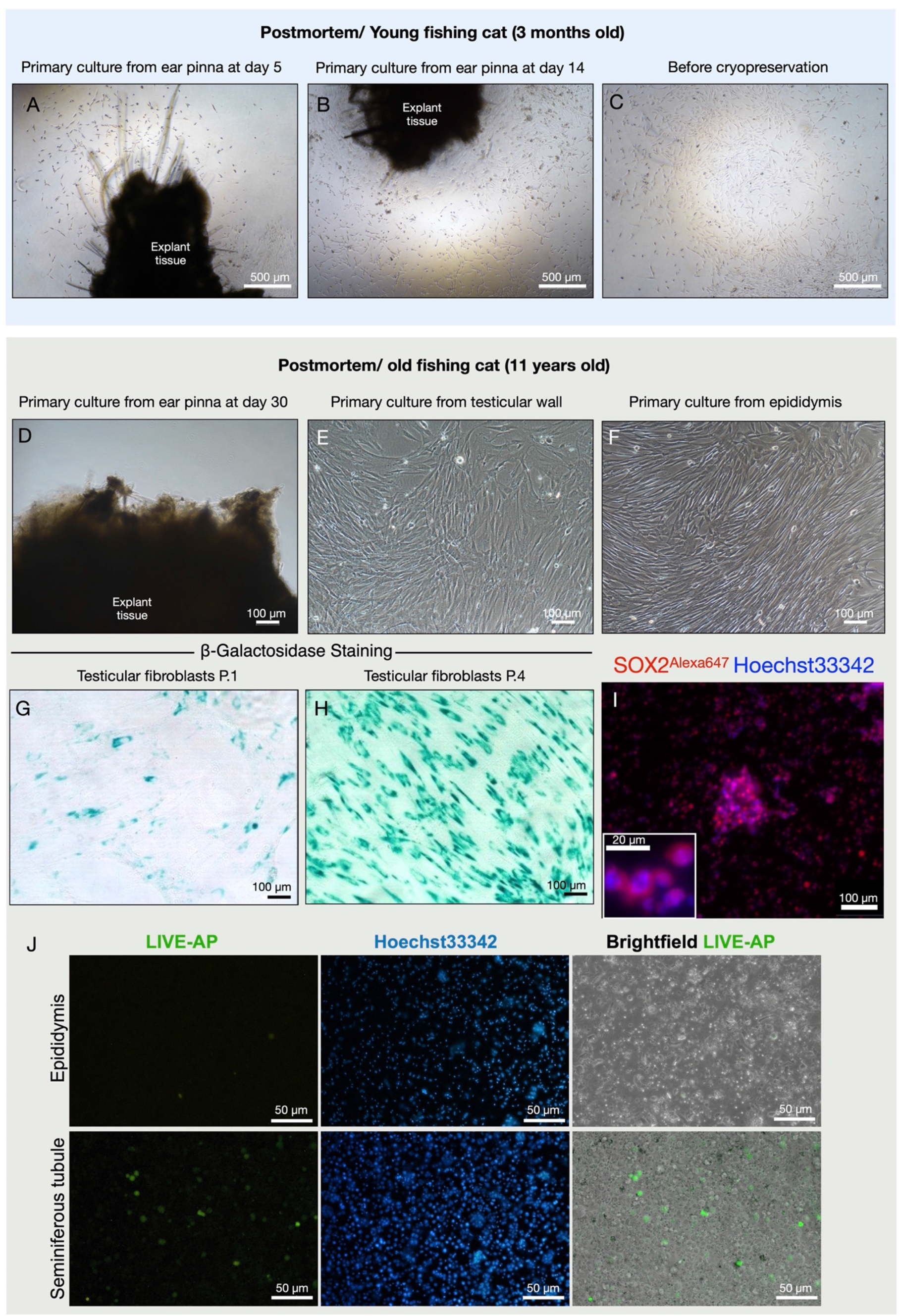
Primary culture of postmortem fishing cats. A-C) Primary culture of tissues collected from young fishing cat at 3 months old. D-J) Primary culture of tissues collected old fishing cat at 11 years old. D) No cell expanded from ear pinna explant in primary culture at day 30. E) Fibroblast outgrowth from testicular wall explants. F) Fibroblast outgrowth from epididymis explants. G-H) Fibroblasts from the testes at passage 1 and passage 4 were tested for cell senescence. I) Immunostaining of seminiferous tubule extract to detect SOX2 protein (Red, Alexa647) with nuclear staining Hoechst33342. Inset shows zoom-up of SOX2 positive cells. J). Live alkaline phosphatase staining was used to detect the presence of putative spermatogonial stem cells (SSCs) from seminiferous tubules and epididymis.

### Delivery of DNA vectors to hard-to-transfect adult fishing cat cells by nucleofection

To enhance limited capacity of fibroblasts to proliferate, here we aimed to optimize DNA delivery into the dermal fibroblasts of fishing cat to further apply for next step of reprogramming. Nucleofection, performed by Nucleofector™ (Lonza), has been used for non-viral transfection of genetic materials with high efficiency into primary cells and hard-to-transfect cells. In this present study, we conducted DNA delivery using 4D-Nucleofector™ system. In general, each cell line requires specific Nucleofector Solution (non-disclosed recipe) and particular program of electric pulse to transfer DNA into cytoplasm and even nucleus of cells. Here we aimed to define optimal Nucleofection™ condition by testing seven programs specific for various types of mammalian cell lines including human cells with recommended Primary Cell 2 (P2) solution specific for mammalian dermal fibroblasts. Based on the expression of GFP, all programs could transfect fishing cat fibroblast cells. Each program provided different cell viability and transfection efficiency. At day 4 post nucleofection when the cells reach 90% confluency, differences in GFP expression became obvious in that three programs including DS-150, EN-150 and FF-135 showed outstanding performance. However, after subculturing, the transfected cells with EN-150 and FF-135 conditions still contained GFP expression (Figure 4). We used flow cytometry to confirm the transfection efficiency and showed that program FF135 can transfect the best with 33.10-41.37% GFP+ cells at day 4 post transfection (Figure 4B). The percentage of GFP+ cells reduced more than half in Day 6 when the cells reach more than 90% confluency (Figure 4C).

**Figure 4.**
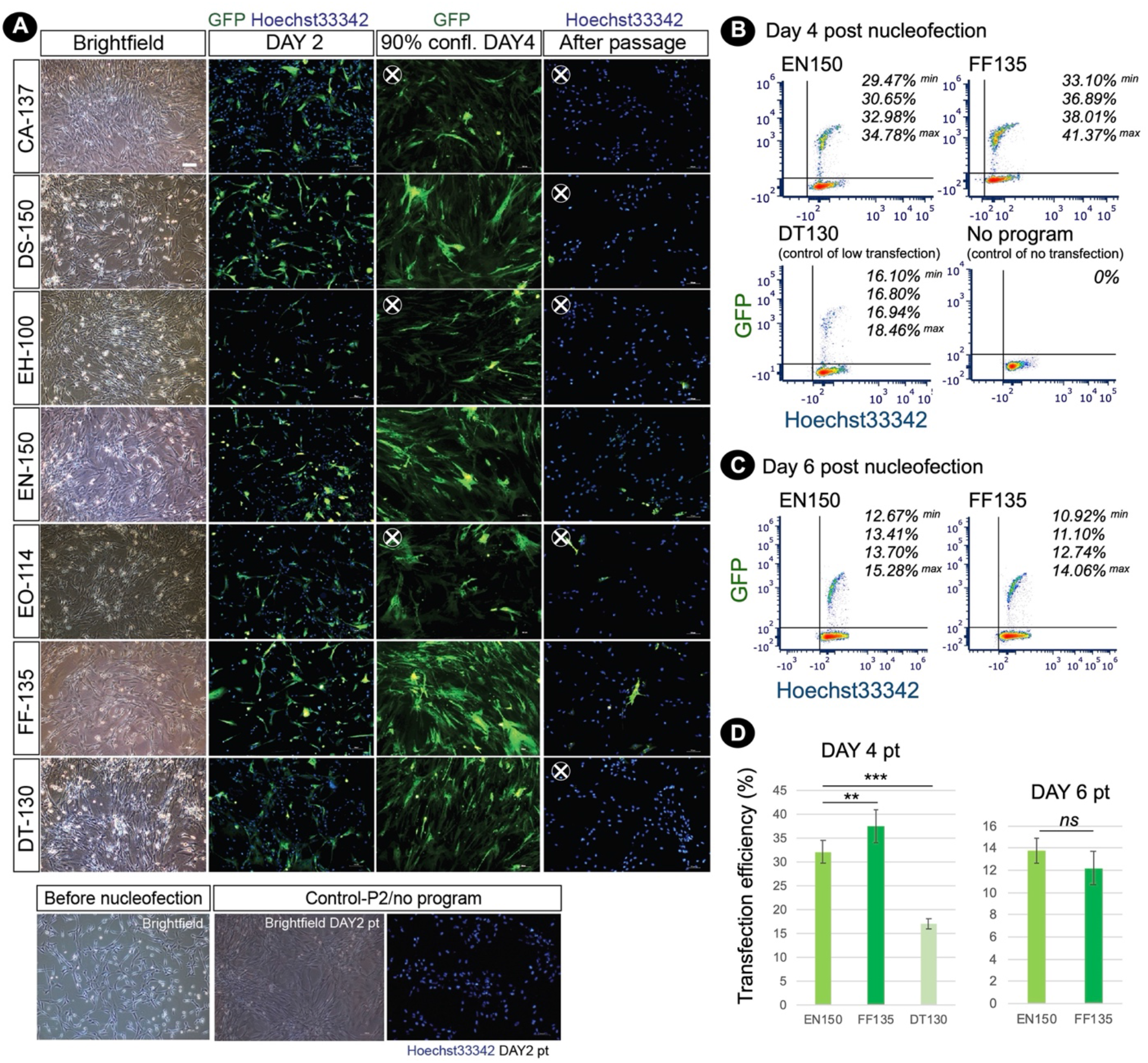
Nucleofection-based DNA delivery into adult dermal fibroblasts from fishing cat. A) Fluorescence photographs show the fibroblasts transfected with pmaxGFP expressing GFP by using nucleofection method. Seven different nucleofection programs (CA-137, DS-150, EH-100, EN-150, EO-114, FF-135 and DT-130) were examined. Photos were taken at day 2, day 4 (90% confluency) and day 6 (day 2 after subculture) post nucleofection. Nuclei were counterstained with Hoechst33342 (Blue). At day 4 post nucleofection, CA-137, EN-150 and EO-114 conditions contained a smaller amount of GFP+ cells and were removed for further analysis (X). After passaging, DS-150 and DT-130 condition shows a smaller number of GFP+ cells and were removed for further analysis (X). B) Cells transfected with the best two nucleofection programs: EN-150 and FF-135 were analyzed by flow cytometer to quantify transfection efficiency. The representative contour plots show the percentage of double GFP+ and Hoechst33342+ cell subsets (4 replicates) at day 4 post transfection. DT-130 program was used as a control of low transfection and “No program” indicates a condition of no nucleofection in the presence of P2 solution and pmaxGFP DNA. C) as in B), the contour plots show the percentage of double GFP+ and Hoechst33342+ cell subsets (4 replicates) at day 6 post transfection or day 2 after subculture. D) Graph shows transfection efficiency of nucleofection (mean ± standard deviation) at day 4 and day 6 post transfection calculated from percentage of double GFP+ and Hoechst33342+ cell subsets (n=4, as shown in B). Asterisks (** and ***) indicate significant difference (p<0.05 and p<0.01 respectively, Student’s t test) and “ns” indicates not significant (p<0.05). Abbreviation: pt, post transfection; confl, confluency.

After nucleofection, some abnormal character of fibroblasts could be detected. Different degree of multinucleated cells appeared in the nucleofected culture within 24 hours. Among the best two Nucleofector™ programs (EN-150 and FF-135) described earlier, FF-135 exhibited higher degree of multinucleation (Figure 5A). However, this multinucleation could be eliminatied by one-time subculture of FF-135 transfected condition (Figure 5B). In addition, to improve survivor of cells after nucleofection, we treated transfected cells with RevitaCell™ Supplement (RC), which contains ROCK inhibitor comparable to Y-27632 and Thiazovivin and found that addition of 0.5X and 1.0X RC improved survived cells in similar manner, which two-fold higher than non-treated transfected cells (Figure 5C).

**Figure 5.**
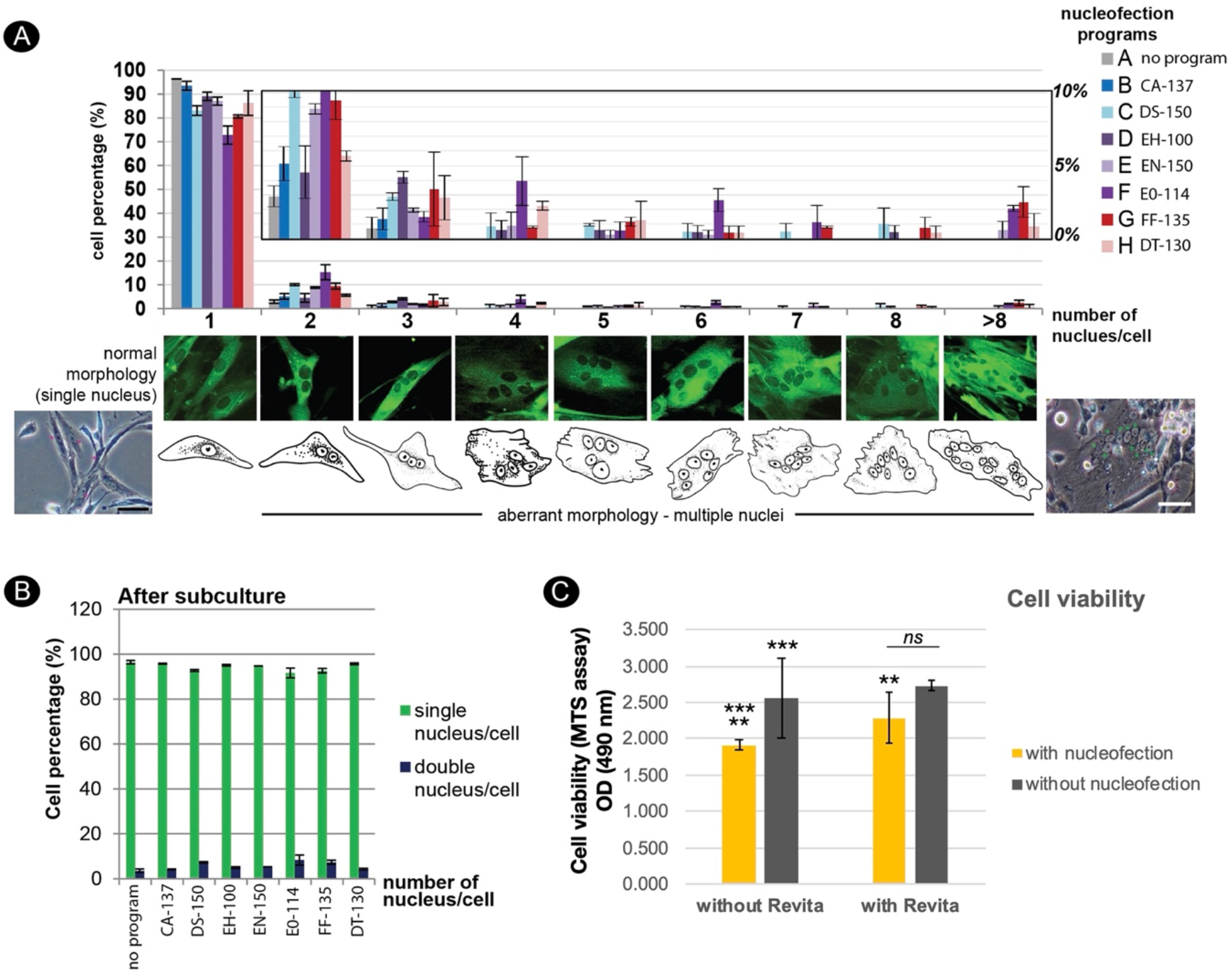
Effect of nucleofection on multinucleation and cell viability of adult dermal fibroblast derived from fishing cat. A) Multinucleation occurred in different levels from various Nucleofector™ programs. Top panel shows a graph depicting incidence of multinucleation from different Nucleofector™ programs. Y axis value represents the percent of transfected cells with one or more nuclei per cell (mean ± standard deviation). X axis indicates the number of nucleus per cell. Middle panel shows morphology of GFP expressing cells with different types of multinucleation. Bottom panel shows brightfield images and cartoons of fibroblast cells with a single nucleus and multiple nuclei. B) Graph depicts number of nucleus per cell as in A) from subculture of the transfected cells at day 3 post passaging or day 7 post transfection. Subculture of transfected cells removed multinucleation from all examined programs. C) Graph shows cell viability of fibroblast cells after nucleofection with program FF-135 with or without RevitaCellTM supplement. Cell viability was done by MTS assay and quantified with spectrophotometer at 490 nm as shown in Y axis. Mean indicates the average of absorbance values from four independent experiments with three technical replicates each. Error bar indicates standard deviation. Asterisks (** and ***) indicate significant difference (p<0.05 and p<0.01 respectively, Student’s t test) and “ns” indicates not significant (p<0.05).

### Challenge to obtain pluripotent stem cells for future conservation

Reprogramming of fishing cat fibroblasts was aimed to enhance cell propagation ability via the induction of pluripotent state. At first, we induced fishing cat fibroblasts with piggyBAC transposon-transposase system by applying nucleofection strategy in previous section with mouse reprogramming factors and mOrange as a reporter (Figure 6A). We can confirm the presence of transfected cells with mOrange expression (Figure 6B) although the iPSC colonies cannot be formed with this approach. In the second strategy of piggyBAC transposon-based reprogramming on fishing cat testicular fibroblasts, we used Tet-on expression system of reprogramming factors and cell selection using puromycin (Figure 6C). The expression of TdTomato allowed us to monitor the expression of reprogramming factors. We found that with this approach iPSC-like colonies with TdTomato expression emerged (Figure 6D). However, TdTomato-expressing colonies was not expandable in iPSC medium with human LIF. We also applied Episomal-based reprogramming using human reprogramming factors (Figure 6E). Episomal vectors-transfected cells were treated with various conditions (Figure 6E). With this approach, fishing cat cells can be expanded the most in FBS-based medium and with Geltrex (Figure 6F). The fibroblastic characters were lost in Geltrex condition, becoming more epithelial characters (Figure 6F), but not in mouse-based feeder cells (Figure 6F). After prolonged culture until the third week of reprogramming, iPSC-like colonies appeared in Geltrex (Figure 6F) but retained flatted epithelial like cells in vitronectin (Figure 6F). However, the picked colonies from Geltrex cannot be propagated. Lastly, we tested whether RNA based reprogramming can induce fishing cat cells based on Yoshioka et al., 2013 and Yoshioka and Dowdy, 2017 [33-34]. We used self-replicating RNA expressing OCT4, SOX2, KLF4, CMYC and GLIS1 proteins to induce the formation of iPSC colonies but the same problems occurred from unexpandable clones (Figure 6G-H).

**Figure 6.**
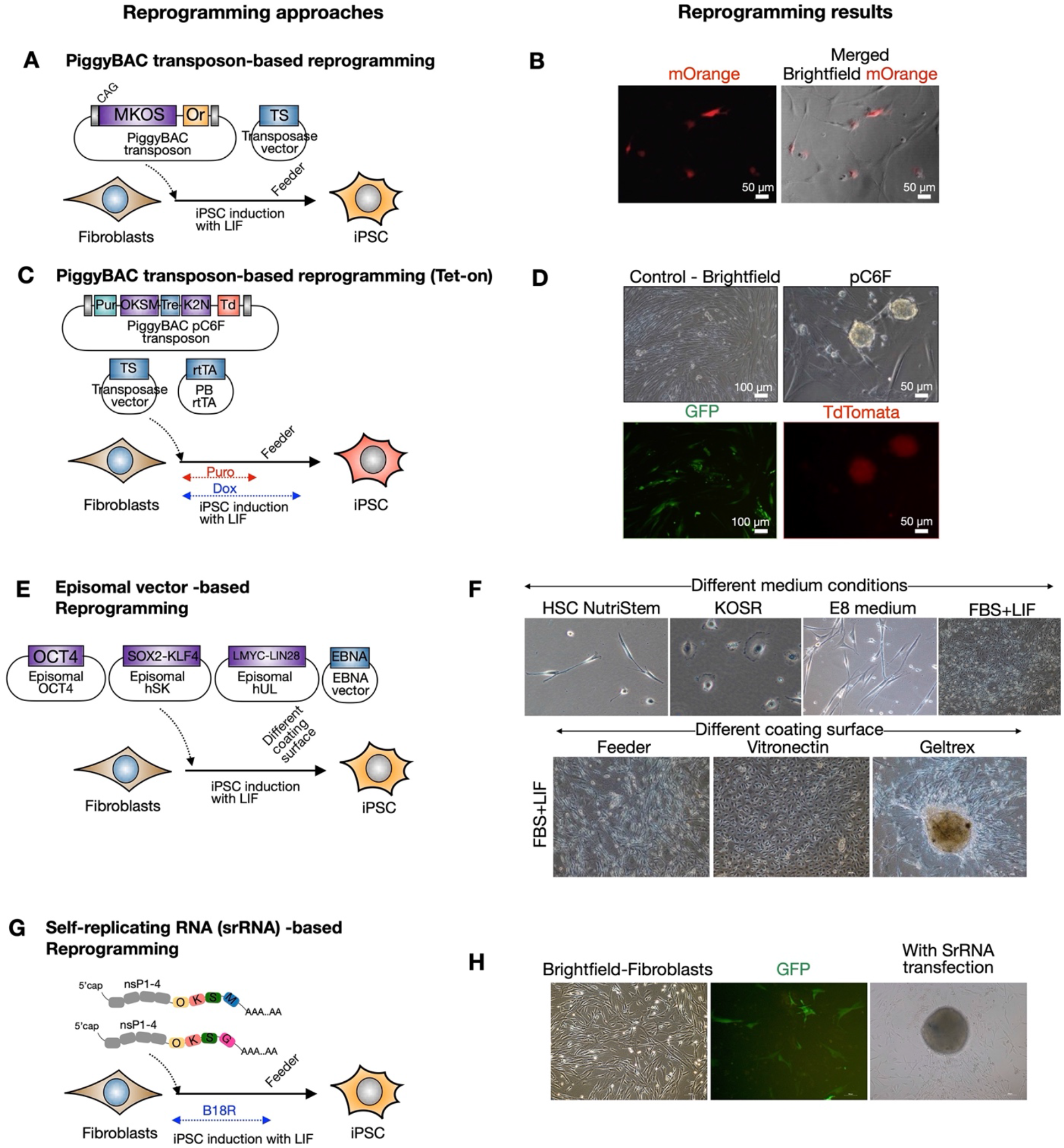
Cellular reprogramming of fishing cat cells via different virus-free approaches. A-B) Fishing cat fibroblasts were reprogrammed with PiggyBAC transposon carrying mouse C-myc, Klf4, Oct4 and Sox2 (MKOS) and mOrange (Or) as a reporter, together with an expression vector expressing transposase (TS). The fibroblasts expressed mOrange, indicating the successful integration of piggyBAC transposon. C-D) The fibroblasts were reprogrammed with piggyBAC transposon carrying inducible reprogramming cassette composed of human OCT4, KLF4, SOX2, C-Myc (OKSM) and human KLF2 and NANOG followed by Ires-Tdtomato. The transfection included transposase vector and piggyBAC expressing rtTA. The transfected cells were selected with puromycin and the expression of reprogramming factors were induced via Tet-on using Doxycycline. The iPSC-like colonies appeared with TdTomata expression. E-F) Reprogramming of fibroblasts using episomal vectors. The transfected cells were tested in various media including commercial HSC NutriStem and E8 medium, KOSR containing medium (KOSR) and media with 10% Fetal Bovine Serum (FBS) with LIF. The induced cells were also replated onto different matrix including mouse irradiated feeder, vitronectin and Geltrex. G-H) RNA-based reprogramming using self-replicating RNA (srRNA) expressing reprogramming factors. The testicular fibroblasts were transfected with srRNA and cultured in the presence of B18R protein before replating onto feeder cells. The iPSC-like colonies appeared in three weeks after transfection.

Thus, inducing and capturing pluripotency of fishing cats is still a main challenge. However, with these reprogramming approaches and adjusted culture strategy, the fishing cat fibroblasts can be reprogrammed toward expandable intermediate cells, in particular with episomal vector based reprogramming (Figure 6), beyond the limitation of fibroblast passaging that we can keep in cryopreservation for future reprogramming success.

## Discussion

Fishing cat cell biobanking is currently in need of preserving its genetic back-up. Here we reported the progress of our cryopreservation of fishing cats from several sources (summarized in Table 1) and provide examples and challenges of using derived adult somatic cells for cellular reprogramming. Embryonic or fetal fibroblasts are common sources for downstream applications as the fibroblasts from embryos exhibited better regeneration process and wound/tendon healing than adult fibroblasts [35-36]. However, embryo sources from wildlife are hard to be obtained. Thus, adult cells are almost only option. In this study, the adult cells can be derived from tissues of both living and postmortem fishing cats but various sources contributed differently, summarized in Figure 7. From the living source, small pieces of ear pinna provided the highest number of fibroblasts for cryopreservation. In contrast, the ear pinna from the postmortem animal contributed to the fibroblast derivation the least. Testis collection from the postmortem fishing cat, on the other hand, provided sufficient cryopreserved sources of cells including fibroblasts from testicular wall and epididymis and mixed cell populations containing SOX2-positive SSCs from seminiferous tubules. Thus, here we recommend collecting testes for cell culture from freshly dead wild animals will provide a good source for biobanking. Although it was not unexpected that keratinocytes appeared from the explant culture of ear pinna carrying intact epidermis, keratinocytes can be eliminated after continuing culture in the medium of DMEM-high glucose with FBS as appeared in Vangipuram et al., 2013 for human fibroblast derivation and Siengdee et al., 2018 for Asian elephant fibroblast derivation [20,37]. In addition to medium types, the procedure of cell culture and supplementation also affected to the culture [20,37]. Either prolonged culture or more passaging of fibroblasts of fishing cats leads to cell senescence and fibroblast is no longer usable for other applications. The obstacles can be relieved by addition of bFGF, a common cytokine known for its ability to support human fibroblast cell proliferation in dose-dependent manner through ERK1/2 and JNK pathways [38], which enhanced cell proliferation in our culture. Hence, frozen stock of early fibroblasts should be made as many as possible, which is also limited to the amount of tissues that can be harvested.

**Figure 7.**
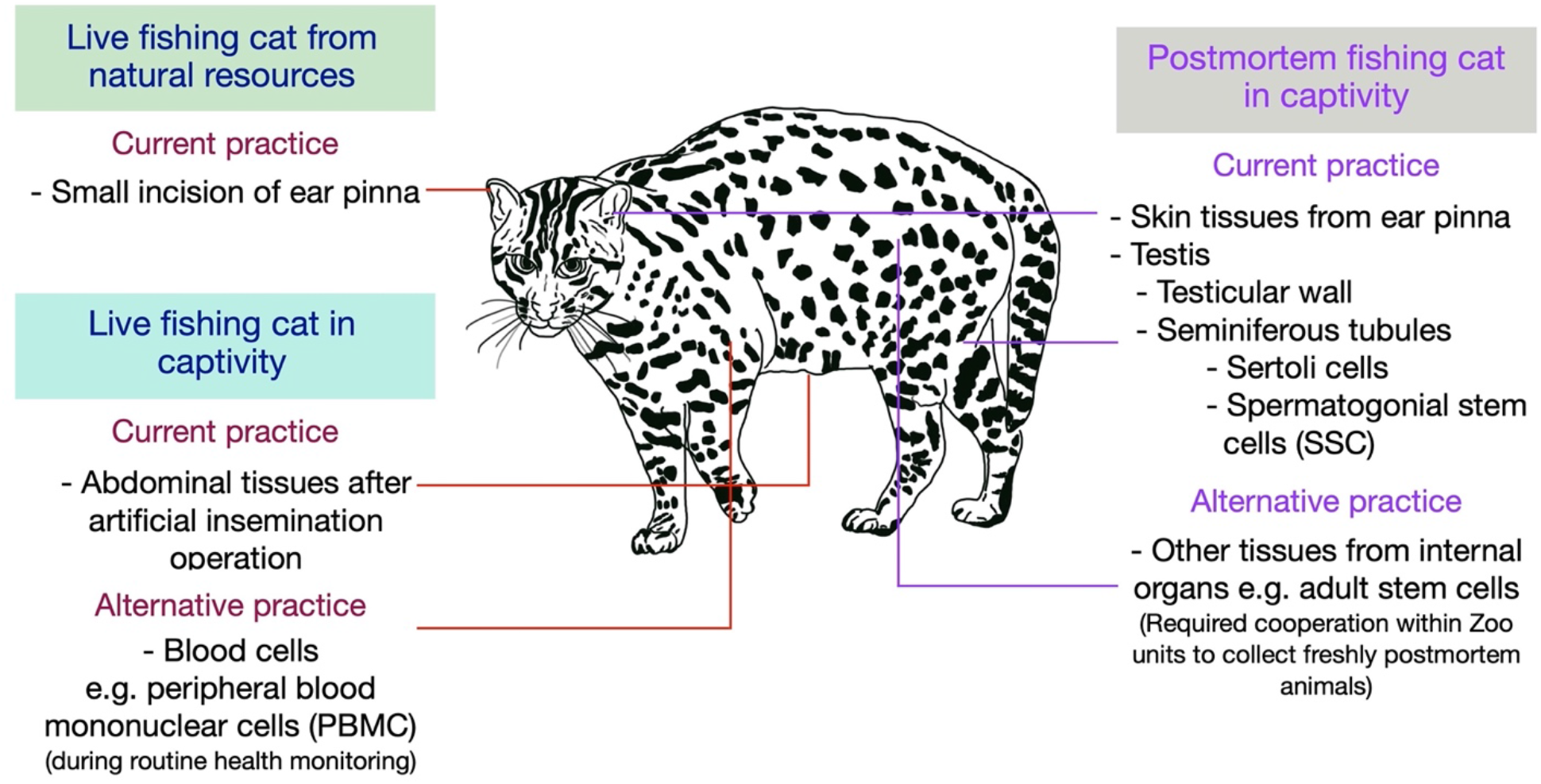
Tissue collection from fishing cat for conservation. Schematic illustration showing various sources of tissue collection for primary cell culture for cryopreservation in this study and suggesting the alternative sources for primary cells from the fishing cats.

In this study, we also show the presence of living SOX2+ spermatogonial cells from postmortem fishing cat. It has been shown that some population of Sox2+ cells, representing a type of adult stem cells in mice, can repopulate the testes with ablated spermatogenesis and restore spermatogenesis [39]. Thus, the presence of SOX2+ cells in extracted seminiferous tubules from the fishing cat open the possibility of adult stem cell derivation and maintenance or even reprogram back to more potential stem cells. Other active adult stem cells in the freshly postmortem animals might be present, which further investigation could help preserving more sources of postmortem wild animals in captivity.

To deliver genetic material to fishing cat adult cells (e.g. for reprogramming) is another challenge due to nature of hard-to-transfect cell type with low division rate compared to embryonic sources. Nucleofection, an electroporation-based transfection method enabling direct DNA delivery into nucleus, can solve the problem as we showed varied range of transfection efficiency from different Nucleofector programs can indeed deliver DNA into the fishing cat cells. Similarly, this technique has been shown to be one of the most suitable non-viral transfection methods to deliver DNA into dermal fibroblasts in various species including human, rat and mouse [28,30-31,40-42].

As the fibroblasts have limited capacity to expand, the preserved cells for better usability is through the reprogramming of the fibroblasts to iPSCs, which have high cell potency to self-renewal and differentiate to all types [43-44]. Felid reprogramming has already been done in some wild cats and domestic cats with retrovirus/lentivirus-based approaches [24,27,45]. Since then there is no success of non-integration approach of wild felid reprogramming. In this study, we examined mouse/human reprogramming system to induce the fishing cat cells without using virus to deliver reprogramming factors. The iPSC colonies appeared from episomal vectors with human OCT4, SOX2, LMYC, KLF4 and LIN28 [46] and piggyBAC transposon with human OCT4, SOX2, KLF4, CMYC, KLF2 and NANOG [32]. However, putative iPSC clones cannot be expanded under mouse iPSC (LIF) or human iPSC maintenance (bFGF) conditions; thus, keeping pluripotent states or inducing the fully reprogrammed state of fishing cat is still a challenge. But the reprogramming to partial state with these reprogramming factors also improved the expandable capacity of the fishing cat cells, contributing more cryopreserved cells for biobanking.

## Conclusion

Generation of biobanking to preserve genetics of fishing cats is limited due to availability of obtaining samples from nature and captivity, fibroblast potency, delivery of genetic material to hard-to-transfect cells and achieving fully reprogramming state. Nevertheless, we succeed in preserving somatic cells from both living and postmortem fishing cats for future conservation technology to prevent the extinction of fishing cats.

## Acknowledgement

We greatly appreciate tremendous efforts of field surveys from Fishing Cat Conservation Project team in Thailand including Supalak Kiatsomboon, Itti Boonorrana, Sudarat Baicharoen, Sriprapai Jumpadang, Saowaphang Sanannu, Waleemas Jairak, Supakorn Pratumratanatarn, Nakorn Slangsingha, Anan Phomphoem, Chaiporn Jaikaeo and Apirak Jansang as well as Panthera Thailand. We thank Wildlife Reproductive Innovation Center, Research Department, Bureau of Conservation and Research and Khoa Kheow Open Zoo, Zoological Park Organization under the Royal Patronage of H.M. the King, Bangkok, Thailand, to provide cell culture and molecular work facilities. We appreciate the technical supports from staffs including Rungwit Chaijirawon, Weerapat Thammasirithong and Kasaraporn Janprasert from Department of Biology, Faculty of Science, Burapha University. We are grateful to Dr. Elke Lorbach, Scientific Support Specialist of Lonza, for helpful information on nucleofection. We thank Dr. Keisuke Kaji for kindly provided PiggyBAC transposon carrying reprogramming factor ORF and reporter mOrange ORF and Dr. Methichit Wattanapanitch for episomal reprogramming approach and materials. We are also grateful for the support from SIF-Science Innovation Facility, Faculty of Science, Burapha University. In addition, we thank the kind support and advice for flow cytometry from Dr.Naphat Chantaravisoot and Dr. Arthid Thim-uam, Center of Excellence in Systems Biology, Faculty of Medicine, Chulalongkorn University.

